# Association of 10 VEGF Family Genes with Alzheimer’s Disease Endophenotypes at Single Cell Resolution

**DOI:** 10.1101/2024.04.12.589221

**Authors:** Yiyang Wu, Julia B Libby, Logan Dumitrescu, Philip L. De Jager, Vilas Menon, Julie A. Schneider, David A. Bennett, Timothy J Hohman

## Abstract

The cell-type specific role of the vascular endothelial growth factors (VEGFs) in the pathogenesis of Alzheimer’s disease (AD) is not well characterized. In this study, we utilized a single-nucleus RNA sequencing dataset from Dorsolateral Prefrontal Cortex (DLFPC) of 424 donors from the Religious Orders Study and Memory and Aging Project (ROS/MAP) to investigate the effect of 10 VEGF genes (*VEGFA, VEGFB, VEGFC, VEGFD, PGF, FLT1, FLT4, KDR, NRP1*, and *NRP2*) on AD endophenotypes. Mean age of death was 89 years, among which 68% were females, and 52% has AD dementia. Negative binomial mixed models were used for differential expression analysis and for association analysis with β-amyloid load, PHF tau tangle density, and both cross-sectional and longitudinal global cognitive function. Intercellular VEGF-associated signaling was profiled using CellChat. We discovered prefrontal cortical *FLT1* expression was upregulated in AD brains in both endothelial and microglial cells. Higher *FLT1* expression was also associated with worse cross-sectional global cognitive function, longitudinal cognitive trajectories, and β-amyloid load. Similarly, higher endothelial *FLT4* expression was associated with more β-amyloid load. In contrast to the receptors, *VEGFB* showed opposing effects on β-amyloid load whereby higher levels in oligodendrocytes was associated with high amyloid burden, while higher levels in inhibitory neurons was associated with lower amyloid burden. Finally, AD cells showed significant reduction in overall VEGF signaling comparing to those from cognitive normal participants. Our results highlight key changes in VEGF receptor expression in endothelial and microglial cells during AD, and the potential protective role of VEGFB in neurons.

## Introduction

Neurovascular unit dysfunction is a hallmark of progression of AD. The vascular endothelial growth factor (VEGF) family of signaling proteins are responsible for the growth and maintenance of cells involved in vascularization and neuralization [1]. The members of the VEGF family include five ligand genes (*VEGFA, VEGFB, VEGFC, VEGFD*, and *PGF*), three receptor genes (*FLT1, KDR*, and *FLT4*), and two co-receptor genes (*NRP1* and *NRP2*; **Figure 1a**). These genes play a role in brain injury [2], stroke [3, 4], and the progression of neurodegenerative disease [5], particularly Alzheimer’s disease (AD). Within AD, disruptions in vascularization and circulation of the brain have been shown to contribute to neurodegeneration and the development of neuropathology [6–13]. As a result, VEGFs have become exciting targets for research to better understand vascular contributions to AD.

**Figure 1.**
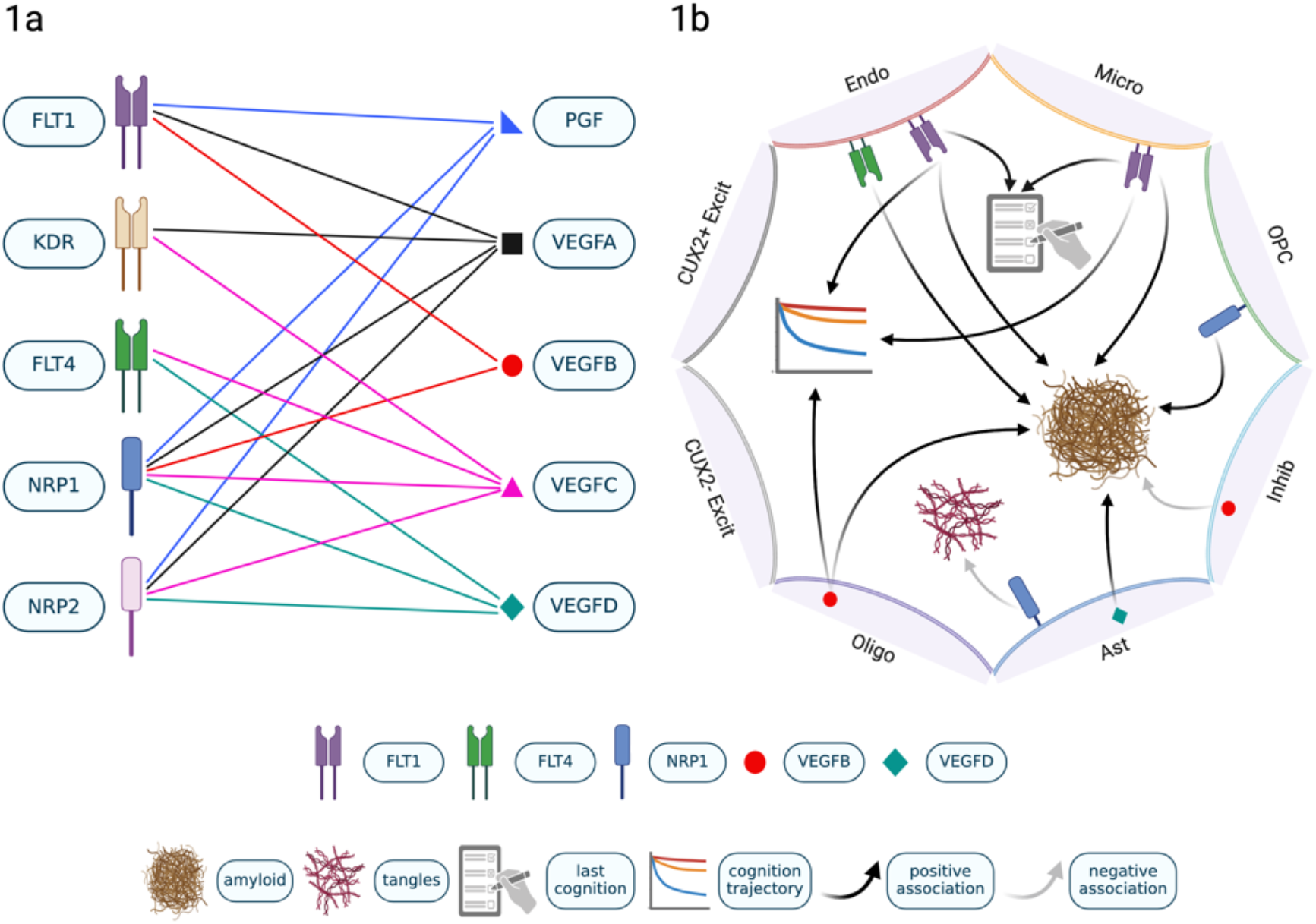
10 VEGF System and Their Cell-Type Specific Effect on AD Endophenotypes 1a. 10 VEGF ligand-receptor system. **1b**. Association of VEGF and AD endophenotypes in DLPFC brain cells. Cell type abbreviations are as follows, Endo, Endothelial Cells; Micro, Microglia; OPC, Oligodendrocyte Precursor Cells; Inhib, Inhibitory Neurons; Ast, Astrocytes; Oligo, Oligodendrocytes; CUX2+ Excit, CUX2+ Excitatory Neurons; CUX2-Excit, all Excitatory Neurons that don’t express CUX2-. This figure was created with BioRender.com.

The role of VEGFs in AD appears to vary depending on the gene family member. Some family members have protective effects [9, 14, 15], while others contribute to the progression of neurodegeneration [16–18]. Earlier work from our group observed that cerebrospinal fluid levels of VEGFA displayed a protective effect [15]. When we expanded our analyses to brain tissues we observed that *VEGFB, FLT1, PGF*, and *FLT4* all appear to contribute to the progression of AD pathology and cognitive decline [16]. Within the prefrontal cortex, higher expression of these four genes (*VEGFB, FLT1, PGF*, and *FLT4*) was associated with more rapid cognitive decline and higher pathology burden [16]. Similar associations are observed when measuring the protein abundance of these VEGF family members in brain tissue and extend to other brain regions including the posterior cingulate gyrus [18].

To better understand VEGF associations in the human brain within their cellular context, we previously investigated single nucleus expression from 48 post-mortem DLPFC samples, evaluating their expression in astrocytes, microglia, oligodendrocytes, oligodendrocyte progenitor cells (OPCs), pericytes, endothelial cells, excitatory and inhibitory neurons [18]. Fascinatingly, we found that the robust *FLT1* and *VEGFB* signals that we had observed in bulk tissue were driven by expression in microglia for both genes, along with endothelial cells, oligodendrocytes, and OPCs for *VEGFB*. While results were encouraging, the small sample size and few endothelial and microglial cells in this initial dataset made interpretation challenging.

The goal of this study is to further disentangle the cell-type specific VEGF expression changes in relation to AD endophenotypes by analyzing a much larger AD single-nucleus RNA sequencing (snRNA) cohort (N=424) [19, 20]. Specifically, we investigated 10 VEGF (*VEGFA, VEGFB, VEGFC, VEGFD, PGF, FLT1, FLT4, KDR, NRP1*, and *NRP2*) family members’ expression within eight brain cell types to test for differential expression between cognitively normal participants and those with AD, along with expression associations with global cognition (cross-sectional and longitudinal trajectories) and AD pathology (β-amyloid and tau neurofibrillary tangle density; **Figure 1b**). In addition, we evaluated cell-type specific changes in the VEGF-associated signaling pathway during AD by conducting an intercellular communication analysis for these 10 VEGF family members.

## Methods

### Participants

The participants included in this study were from two longitudinal clinical-pathological cohort studies including the Religious Orders Study and the Rush Memory and Aging Project (ROS/MAP). Each study was approved by an Institutional Review Board (IRB) of Rush University Medical Center. At enrollment, all participants were free of known dementia and agreed to annual clinical evaluation and the donation of their brain at the time of their death [21–23]. All participants signed informed and repository consents and an Anatomic Gift Act. Secondary analyses of this extant data were approved by the Vanderbilt University Medical Center IRB. ROS/MAP data can be accessed online at the Rush Alzheimer’s Disease Center Resource Sharing Hub (https://www.radc.rush.edu/), as well as on the Accelerating Medicines Partnership-Alzheimer’s Disease (AMP-AD) Knowledge Portal (syn3219045).

### Single nucleus RNA sequencing

Specimens were collected on availability of frozen pathologic materials from the dorsal lateral prefrontal cortex (DLPFC). The tissue specimens were collected by the Rush Alzheimer’s Disease Center and processed at Columbia University Medical Center. Participants with a post-mortem interval < 41 hours and whole genome sequence data were selected to go through a series of rigorous quality control steps [24–26], which resulting in 424 participants. The participants had a median of 3,824 sequenced nuclei. In the final data, there were 8 major cell types, including astrocytes, endothelial cells, inhibitory neurons, microglia, oligodendrocytes, oligodendrocyte precursor cells (OPCs), and CUX2- and CUX2+ excitatory neurons. snRNAseq data can be downloaded from AD Knowledge Portal (Accession Number: syn31512863).

Additional filters applied before analyses included removing genes with expression in less than 10% of all cells, removing cells with less than 200 or more than 20,000 total RNA UMIs or had more than 5% mitochondrial mapped reads.

### Measurements of Cognitive Function

The measurement of global cognition was derived from 17 different neuropsychological tests across five domains of cognition (semantic, episodic, and working memory, perceptual speed, and perceptual orientation). The z-scores of all the available tests were averaged to create a global cognition composite. Further details on the development of this composite have been previously described [27]. For cross-sectional cognition function related analysis, global cognitive score at last visit before death was used from 369 participants. Longitudinal cognitive trajectory was derived from a linear mixed effects model with global cognition as the outcome and the intercept and interval (years from last visit) entered as fixed and random effects. The derived “cognition slope” was calculated as the sum of the random and fixed effects for each participant. 423 participants are included in this analysis.

### Measurements of AD pathology

Measures of pathology were previously characterized in ROS/MAP [21, 22]. Measured by immunohistochemistry at autopsy, β-amyloid load and tau tangle density were quantified as the average percent area occupied by β-amyloid (using antibodies specific to Aβ_42_) or tau (using antibodies specific to AT8 epitope of abnormally phosphorylated tau) across 8 brain regions at autopsy: hippocampus, angular gyrus, and entorhinal, midfrontal, inferior temporal, calcarine, anterior cingulate, and superior frontal cortices. Then, values were transformed to approximate a normal distribution. 421 participants are included in β-amyloid load and tau density analysis.

### Statistical Analysis

Statistical analyses were completed using RStudio (R version 4.3.1) and code is available upon request from the corresponding author. Significance was set to a priori to α=0.05.

Additionally, p-values were corrected for all *VEGF* predictors across AD-associated outcomes (290 models) using the false discovery rate (FDR) procedure.

Negative binomial mixed models implemented by the Nebula R package [28](v1.4.2) were used to analyze the association between each *VEGF* gene within each of the eight cell types and AD diagnosis, global cognition longitudinally and at last visit before death, and β-amyloid load and PHF tau tangle density at death. For analysis of diagnosis, 299 individuals were included: 142 normal cognition controls (cogdx = 1) and 157 AD dementia patients (cogdx = 4 or 5). All models covaried for age at death, sex, post-mortem interval (PMI), and the interval between last cognitive visit and death in years (for cognition). One participant didn’t have PMI data, hence, was removed from association analysis. The gene count matrix input of the models was the UMI count data from the RNA assay normalized and scaled by the “sctransform” R package (https://github.com/satijalab/sctransform).

### VEGF-mediated Intercellular Communication Profiling

Intercellular communication pattern involving these 10 VEGF genes among cell types was analyzed using the ‘CellChat’ R package [29](v1.0). Due to lack of some of the evident ligand-receptor pairs involving these 10 VEGF genes from the default CellChat database (CellChatDB), we manually updated CellChatDB by incorporating those curated additionally from the CellTalk DB (v1.0, https://github.com/ZJUFanLab/CellTalkDB/tree/master/database). All VEGF ligand-receptor pairs that were included in this study can be found in **Supplemental Table 2**. We performed the rest of the analysis according to the default setting of CellChat, skipping projecting gene expression data onto human protein-protein interaction network.

## Results

### Participant Characteristics

The snRNAseq data used for our analysis consisted of 424 non-Latinx White participants (**Table 1**) with an average PMI of 7.7 hours (SD = 5.1 hours), 68% of whom are females, 52% are clinical AD dementia patients (cogdx = 4 or 5), and 27% carried at least one APOE4 allele. The cohort has a mean age at death at 89 years old with an average of 13 years of education.

**Table 1.** Characteristics of Study Participants.

|  | **Cognitive**<br>**Normal Controls**<br>**(N=143)** | **AD Patients**<br>**(N=157)** | **Total Cohort**<br>**(N=424)** | **P Value** |
| --- | --- | --- | --- | --- |
| Age at death<br>(years) | 87.20 ± 7.26 | 91.15 ± 6.56 | 89.19 ± 6.82 | <0.001 |
| Education<br>(years) | 16.28 ± 3.45 | 16.55 ± 3.22 | 16.31 ± 3.47 | 0.48 |
| Post-mortem<br>interval (hours) | 7.80 ± 4.99 | 7.25 ± 4.32 | 7.71 ± 5.11 | 0.31 |
| Global cognition<br>slope | -0.04 ± 0.04 | -0.19 ± 0.09 | -0.11 ± 0.09 | <0.001 |
| β-amyloid load | 3.12 ± 3.77 | 5.93 ± 4.46 | 4.46 ± 4.50 | <0.001 |
| Tau density | 2.97 ± 3.36 | 9.36 ± 9.15 | 5.98 ± 7.15 | <0.001 |
| Male, no. (%) | 49 (34) | 42 (27) | 136 (32) | 0.16 |
| APOE4 carrier,<br>no. (%) | 27 (19) | 60 (38) | 109 (27) | <0.001 |

Values are presented as mean ± standard deviation unless otherwise indicated. P values of comparison between cognitive normal controls and AD patients were derived from Welch Two Sample t-test for continuous variables and Pearson's Chi-squared test for categorical variables.

### *VEGFs* Associations with Cognition

Higher expression of *FLT1* in endothelial cells and microglia was associated with worse cognitive performance at last visit before death (endothelial-*FLT1* (logFC= -0.089, FDR= 0.009) and microglia-*FLT1* (logFC= -0.176, FDR= 0.009)) and faster cognitive decline (endothelial-*FLT1* (logFC= -0.771, FDR= 0.041) and microglia-*FLT1* (logFC= -1.964, p= 0.009)). Additionally, higher expression of *VEGFB* in oligodendrocytes (logFC= -0.561, FDR= 0.019) was also associated with faster cognitive decline. **Table 2** listed all statistically significant associations between 10 VEGF gene expression and various AD outcomes tested (the complete result can be found in **Supplemental Table 1**).

**Table 2.** FDR Significant Cell-Type Specific VEGF Associations with Cognition and AD Pathology.

| **Gene** | **Outcome** | **Cell Type** | **logFC** | **SE** | **P Value** | **FDR** |
| --- | --- | --- | --- | --- | --- | --- |
| *FLT1* | Last cognition before death | Endothelial cells | -0.089 | 0.024 | 1.768E-04 | 0.009 |
| *FLT1* | Last cognition before death | Microglia | -0.176 | 0.046 | 1.492E-04 | 0.009 |
| *FLT1* | Cognition trajectory | Endothelial cells | -0.771 | 0.248 | 0.002 | 0.041 |
| *FLT1* | Cognition trajectory | Microglia | -1.964 | 0.504 | 9.716E-05 | 0.009 |
| *VEGFB* | Cognition trajectory | Oligodendrocytes | -0.561 | 0.166 | 7.232E-04 | 0.019 |
| *FLT1* | Diagnosis | Endothelial cells | 0.195* | 0.055 | 4.232E-04 | 0.014 |
| *FLT1* | Diagnosis | Microglia | 0.452* | 0.113 | 6.378E-05 | 0.009 |
| *FLT1* | β-amyloid load | Microglia | 0.053 | 0.011 | 2.700E-06 | 7.830E-04 |
| *VEGFB* | β-amyloid load | Oligodendrocytes | 0.014 | 0.004 | 1.428E-04 | 0.009 |
| *VEGFB* | β-amyloid load | Inhibitory neurons | -0.012 | 0.004 | 6.529E-04 | 0.019 |
| *FLT4* | β-amyloid load | Endothelial cells | 0.027 | 0.008 | 3.579E-04 | 0.013 |
| *VEGFD* | β-amyloid load | Astrocytes | 0.016 | 0.005 | 0.001 | 0.029 |
| *NRP1* | β-amyloid load | Oligodendrocyte precursor cells | 0.018 | 0.006 | 0.002 | 0.046 |
| *NRP1* | Tau density | Astrocytes | -0.014 | 0.004 | 2.229E-04 | 0.009 |

\*logFC values of diagnosis is log2 fold change of gene expression comparing AD to cognitive
normal participants.

### *VEGFs* Associations with AD Pathology

Multiple cell types and *VEGF* genes were associated with β-amyloid load (**Table 2**). Higher microglia-*FLT1* (logFC= 0.053, FDR= 0.001), oligodendrocyte-*VEGFB* (logFC= 0.014, FDR= 0.009), endothelial-*FLT4* (logFC= 0.027, FDR= 0.013), astrocyte-*VEGFD* (logFC= 0.016, FDR= 0.029), and oligodendrocyte precursor cell-*NRP1* (logFC= 0.018, FDR= 0.046) were all associated with higher β-amyloid load. On the contrary, within inhibitory neurons, higher expression of *VEGFB* was associated with lower β-amyloid load (logFC= -0.012, FDR= 0.019). On the other hand, we only found one gene that associated with tau density, which was higher astrocyte-expressed NRP1 was associated with lower tau density (logFC= -0.014, FDR= 0.009, **Table 2**).

To examine whether contribution of micro-*FLT1* and oligo-*VEGFB* to more amyloid load influences the association of their expression with worse cognition performance, we performed a competitive hierarchical analysis (“gene expression ∼ sex + age at death + PMI + amyloid + cognition (either cross sectional score at last visit or longitudinal trajectory)” using the same Nebula setting for other association analyses). We found that when covarying for amyloid, both gene associations with cognitive performance were attenuated about 20% (by comparison of logFC values), but remained statistically significant for micro-*FLT1* (p = 0.011 for both cross-sectional and longitudinal cognitive performance models) and fell just below statistical significance in the case of oligo-*VEGFB* (p = 0.065 for longitudinal cognitive performance model). Together our analysis indicated that the associations of micro-*FLT1* and oligo-*VEGFB* to cognitive performance are only partially explained by their association with amyloid load.

### Differential Expression of *VEGFs in AD*

We also tested for differences in *VEGF* expression between neuropathologically confirmed AD dementia cases and cognitively unimpaired. We observed that AD patients had higher expression levels of *FLT1* within microglia (logFC= 0.452, FDR= 0.009) and endothelial cells (**Table 2**, logFC= 0.195, FDR= 0.014).

### VEGF-associated Intercellular Communication Profile

Overall VEGF communication strength (calculated by summing communication probabilities of all curated VEGF pathway ligand-receptor (L-R) pairs among all cell types (**Supplemental Table 2**)) was significantly reduced (Wilcoxon test; p<0.05) in the AD group compared to the cognitively normal group, and the reduction was observed for both incoming and outgoing VEGF signaling strength (calculated by in-degree and out-degree network centrality scores respectively) among cells (**Figure 2a**). VEGFA was the most abundantly expressed VEGF ligand among investigated, primarily in astrocytes, hence, it was no surprise to see many of the significant interactions (L-R pairs) for VEGF pathway were with VEGFA for both AD and cognitively normal groups. As a result, overall VEGF information flow among these brain cells was from astrocytes to other cell types including themselves (**Figure 2b**). But interestingly, the top receiver (cell type) of VEGF communications for the cognitively normal group was excitatory neurons, whereas for AD group, it was endothelial cells (**Figure 2c**). The underlying cause of this difference lay in the significant change of the relative contribution of top L-R pairs to the overall VEGF communications when comparing these two diagnoses groups. Specifically, “VEGFA-GRIN2B” was 37.8% in AD vs 39.5% in cognitive normal group, “VEGFA-SIRPA” was 10.2% vs 10.7% (AD vs cognitive normal), and “VEGFA-EGFR” was 10.2% vs 9.5% (AD vs cognitive normal). But probably the most important change is for “VEGFA-FLT1”, which contributed 13.8% of all VEGF signals in AD dementia patients comparing to 9.5% in cognitive normal individuals. VEGFA-FLT1 signal is from astrocytes to endothelial cells exclusively, and it showed a comparable communication probability between cells in AD and cognitive normal participants (adjusted probability of 0.073 vs 0.076), which can be seen as an outsider considering the overall reduction of VEGF signals in AD group (**Supplemental Table 3** presented the complete result). The significant elevated *FLT1* expression in endothelial cells from AD patients might have facilitated this change.

**Figure 2.**
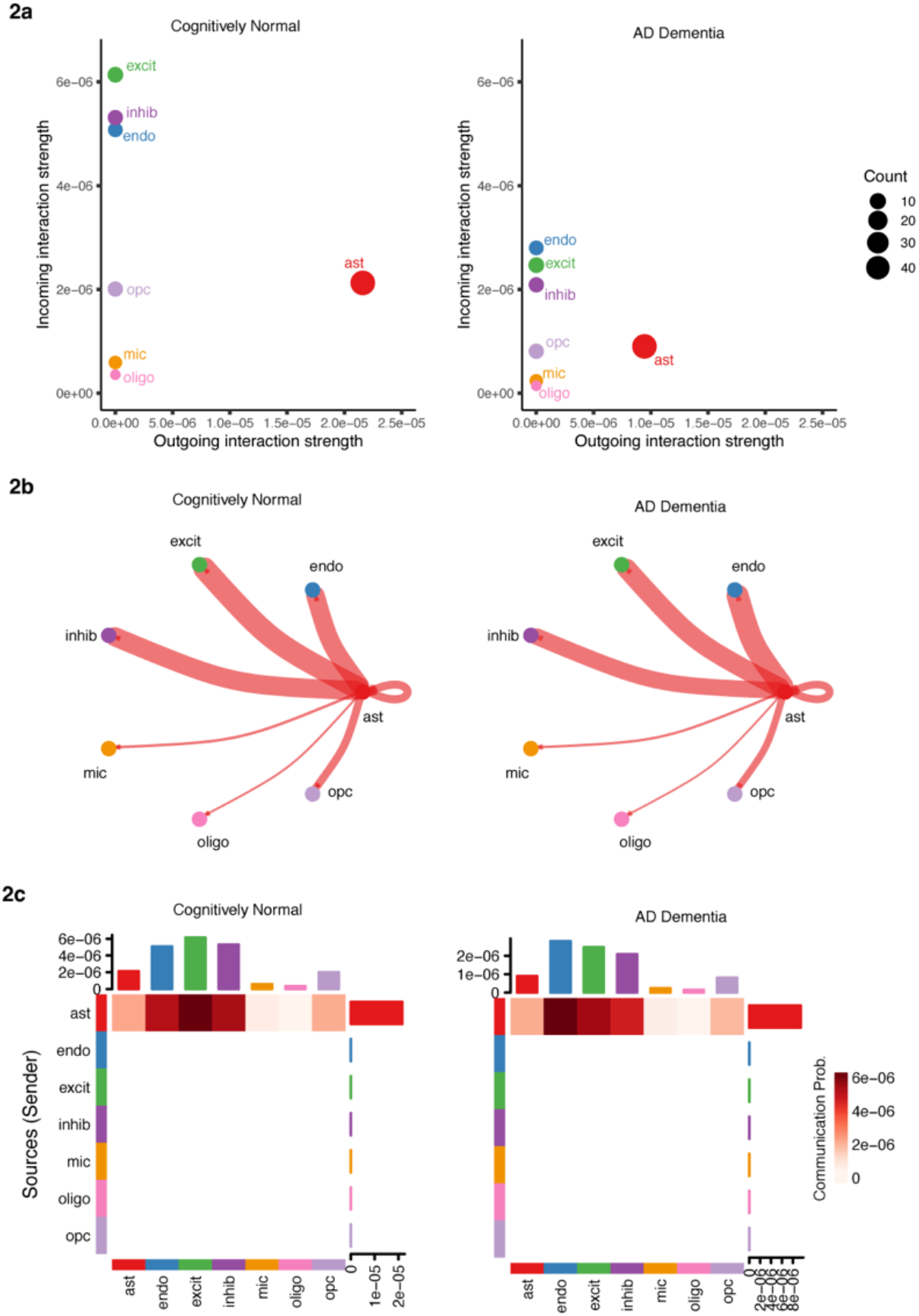
10 VEGFs-Associated Signaling Pathway Profile in AD vs Cognitively Normal Groups 2a. Overall incoming and outgoing VEGF signaling strength comparison. **2b**. Circle plot of overall VEGF information flow. **2c**. heatmap of overall VEGF information flow. Cell type abbreviations are as follows, Endo, Endothelial Cells; Micro, Microglia; OPC, Oligodendrocyte Precursor Cells; Inhib, Inhibitory Neurons; Ast, Astrocytes; Oligo, Oligodendrocytes; CUX2+ Excit, CUX2+ Excitatory Neurons; CUX2-Excit, all Excitatory Neurons that don’t express CUX2-.

## Discussion

In this study we provided the most comprehensive exploration into cell-specific effects of the VEGF family in the context of AD. Our results build on previous evidence from bulk tissue at the protein and transcriptomic level that *VEGFB, FLT1, FLT4*, and *NRP1* relate to the clinical progression and neuropathology of AD [16, 18]. The single cell resolution here builds on our early single nucleus data [18] to provide the clearest picture of the important and distinct roles the VEGF family members play in various brain cell types. Our results particularly highlight the diverse associations for VEGF family members across endothelial cells, microglial cells, astrocytes, oligodendrocytes, and neurons. While *FLT1* showed strong upregulation in the AD brain in endothelial cells as might be expected, we also observed strong upregulation of *FLT1* in microglial cells. Yet, regardless of the upregulation of *FLT1*, our intercellular communication analyses suggest that FLT1-associated canonical VEGF signaling pathways themselves are in fact comparable in the AD brain to normal brain. In contrast to the consistent effects observed for *FLT1*, we observed opposing direction of effect on β-amyloid load for *VEGFB* expression in oligodendrocytes vs inhibitory neurons, whereby higher levels of oligodendrocyte *VEGFB* related to higher pathology, while higher levels of neuronal *VEGFB* related to lower pathology. Together, these results suggest that in addition to the classic vascular signaling pathway underlying the associations between VEGF family members and AD, there may also be immune (e.g., microglial *FLT1*) and neuronal (e.g., neuronal *VEGFB*) signaling cascades that contribute to AD pathogenesis.

The central role of VEGF signaling is angiogenesis under both health and disease conditions [30, 31]. Pathological angiogenesis is a hallmark of AD, which is believed to both drive and respond to multiple AD pathologies driving hypoxia, oxidative stress, inflammation, and Aβ accumulation [32]. Indeed, previous work has identified a marked upregulation of angiogenic factors and mediators in the AD brain [33, 34]. In the present results, the upregulation of *FLT1* in endothelial cells among AD patients could reflect a repair response. The increase in vascularization and blood flow could be a response to AD neuropathology similar to the response that has been indicated in other neurodegenerative and neuropsychiatric diseases such as ALS [35] and schizophrenia [36]. On the other hand, the elevated endothelial *FLT1* could drive AD neuropathology. For example, it has been shown that VEGF-induced pathological angiogenesis can facilitate Aβ generation [37]. VEGF-induced pathological angiogenesis can also worsen AD pathology by disrupting permeability of the blood-brain barrier (BBB) [38, 39]. BBB is formed by physical interaction of multiple brain cell types, with a common structure consisting of blood capillaries formed by endothelial cells, pericytes, and the astrocytic end feet [40]. As a natural protective barrier of the central nervous system (CNS), BBB homeostasis is strictly regulated, disruption of which can lead to many neurological conditions including AD [41, 42]. Our intercellular communication profile showed that signals communicated among these 10 VEGF genes were predominantly from astrocytes to endothelial cells, hinting at the canonical angiogenesis pathway at the blood brain barrier as a primary driver here. In fact, evidence is already emerging that demonstrates CNS angiogenesis can be regulated through VEGF-dependent signaling at the BBB [43]. Characterizing the interplay between VEGF family genes and BBB integrity in the context of AD may provide more insight into the mechanistic pathways underlying the associations observed here.

Our results also suggest roles of some of the VEGF family genes in regulating the immune response, including the upregulation of microglial *FLT1* in AD group. Increased microglia function in AD has been observed by many [44–48] in the presence of β-amyloid peptides and facilitates an inflammatory response [49]. FLT1 was believed to be one of the 20 or so proteins that has been found to be enriched in Aβ plaques [50], and has recently been showed to interact with VCAM1 and plays a role in VCAM1-dependent chemotaxis of microglia, which enhances microglia clearance of Aβ [51]. Studies have also shown the critical role that FLT1 plays in microglial proliferation and monocyte-macrophage migration [52, 53]. With all these under consideration, elevation of microglial *FLT1* in AD should not be a surprise. The primary ligand for *FLT1* is *VEGFB*, so it is possible that the *VEGFB* association in oligodendrocytes may reflect upregulation of this same pathway. It is well known that oligodendrocytes and microglia work together during regulation of myelination, and this is particularly noticeable during neurodegenerative conditions [54]. Future work confirming these cell-specific transcriptomic effects at the protein level is necessary.

One fascinating discovery in the present results is that higher expression of *VEGFB* within inhibitory neurons was associated with lower β-amyloid load, which was the only protective effect of any VEGF we tested in this dataset. This result is consistent with the previously reported evidence of the benefits of *VEGFB* expression on neuronal survival and nerve regeneration [55–58]. Perhaps in the early stage of AD when β-amyloid load is low, inhibitory neurons manage to inhibit apoptosis and promote neurogenesis through upregulating VEGFB. Upregulation of VEGFB could be one of the molecular mechanisms that enables inhibitory neurons to be more resistant to AD neuropathology comparing to excitatory neurons [59]. Interestingly, a study of APP/PS1 amyloidopathy mice did show an increase of inhibitory synapses in hippocampus before pathology develops and then decreased at 12 months of age when robust AD pathology was present [60]. Hence, the protective effect of VEGFB from inhibitory neurons to the development of AD pathology could function in a time-sensitive matter. One also cannot rule out the possibility that increased *VEGFB* expression within inhibitory neurons could help with the clearance of β-amyloid.

We also observed higher expression of *FLT4* within endothelial cells related to higher β-amyloid load. The primary ligand for *FLT4* is *VEGFD* and we also observed an association between elevated *VEGFD* in oligodendrocytes and β-amyloid load. In contrast to the other VEGF receptors, the FLT4*/*VEGFD signaling axis is primarily involved in lymphangiogenesis. Certainly meningeal lymphatics has been implicated in β-amyloid clearance through a complex interplay with the immune system [61], so it is quite possible that lymphatics underly the observed *VEGFD* and *FLT4* associations with β-amyloid. In previous work, we identified an associated between higher expression of *FLT4* in the dorsal lateral prefrontal cortex and worse neuropathology [16], but also identified higher expression of *FLT4* in the caudate nucleus associated with lower β-amyloid levels and tau levels [18]. Others have also observed these flips in direction with associations depending on the brain region [62]. Combined, these results suggest that tissue and cell-type are important considerations when looking at the role of *FLT4* in the brain within the context of AD.

Finally, we observed a cell-type specific effect of NRP1. Higher expression of *NRP1* in astrocytes was associated with lower levels of neurofibrillary tangles, but higher expression of *NRP1* within oligodendrocyte precursor cells was associated with higher β-amyloid load. Other work from our group has also observed differing roles of *NRP1* when pertaining to AD neuropathology [18]. However, this is the first time we have observed statistically significant associations within specific cell-types. All above results further highlight the complexity of the *VEGF* family and their role within neurodegeneration. To fully understand the clinical potential of *VEGF* as a biomarker and therapeutic target, in-depth understanding of each of the tissue and cell-type is necessary.

This study has many strengths. ROS/MAP is a well-characterized cohort with longitudinal cognitive data, comprehensive neuropathology data with multiple cell-type measures of *VEGF* expression. Additionally, this study reports novel discoveries and replication of the effects of *VEGF* family members on AD phenotypes including pathology and cognition at the transcriptomic level within a large sample size. Because of the larges sample size and cell-type specific expression measurements, we were able to provide more comprehensive data and hypotheses about cellular context.

Despite these strengths, this study also has limitations. ROS/MAP is enriched for non-Hispanic white, highly educated individuals which limits the generalizability to more representative populations. Further, while being able to examine cell-type specific effects in a large sample size, we are limited to the dorsolateral prefrontal cortex tissue.

To conclude, this study provides an extensive characterization within a large sample size of the *VEGF* signaling family members in the context of cognitive aging and AD. Our results provide evidence that further supports the role of *VEGFB, VEGFD, FLT1, FLT4* and *NRP1* play inside and outside of classic VEGF signaling axis in AD, with support for both an endothelial and microglial signaling cascade that could underly these well-established associations. For the first time, we provide evidence that *VEGFB* expression in inhibitory neurons may relate to a protective effect against AD neuropathology as has been hypothesized previously and observed in models of Parkinson’s disease. These results provide compelling evidence that VEGFB and FLT1 may have potential as therapeutic targets in AD, but there remains a need to confirm these single nucleus transcriptomic results at the protein level and establish causality through experimental approaches.

## Acknowledgements

This project is supported by R01-AG061518, R01-AG074012, and R01-AG059716. ROSMAP is supported by P30AG10161, P30AG72975, R01AG15819, R01AG17917, U01AG46152, and U01AG61356. ROSMAP resources can be requested at https://www.radc.rush.edu and www.synpase.org.

## AI Content Disclaimer

None of the content used in this article was generated by AI.

## References

1. de Almodovar CR, Lambrechts D, Mazzone M, Carmeliet P. Role and therapeutic potential of VEGF in the nervous system. Physiol Rev. 2009;89:607–648.

2. Linton AE. Pathologic sequelae of vascular cognitive impairment and dementia sheds light on potential targets for intervention. Cereb Circ-Cogn Behav. 2021;2:100030.

3. Greenberg DA. Vascular Endothelial Growth Factors (VEGFs) and Stroke. Cell Mol Life Sci. 2013;70:1753–1761.

4. Ng TKS, Ho CSH, Tam WWS, Kua EH, Ho RC-M. Decreased Serum Brain-Derived Neurotrophic Factor (BDNF) Levels in Patients with Alzheimer’s Disease (AD): A Systematic Review and Meta-Analysis. Int J Mol Sci. 2019;20:257.

5. Melincovici CS. Vascular endothelial growth factor (VEGF) – key factor in normal and pathological angiogenesis. Rom J Morphol Embryol. 2018;59:455–467.

6. Silva T. Circulating levels of vascular endothelial growth factor in patients with Alzheimer’s disease: A case-control study. Behav Brain Res. 2023;437.

7. Ali M. VEGF Paradoxically Reduces Cerebral Blood Flow in Alzheimer’s Disease Mice. Neurosci Insights. 2022;17.

8. Guo L-H, Alexopoulos P, Perneczky R. Heart-type fatty acid binding protein and vascular endothelial growth factor: cerebrospinal fluid biomarker candidates for Alzheimer’s disease. Eur Arch Psychiatry Clin Neurosci. 2013;263:553–560.

9. Huang L, Jia J, Liu R. Decreased serum levels of the angiogenic factors VEGF and TGF-β1 in Alzheimer’s disease and amnestic mild cognitive impairment. Neurosci Lett. 2013;550:60–63.

10. Harris R, Miners JS, Allen S, Love S. VEGFR1 and VEGFR2 in Alzheimer’s Disease. J Alzheimers Dis. 2018:1–12.

11. Provias J, Jeynes B. Reduction in Vascular Endothelial Growth Factor Expression in the Superior Temporal, Hippocampal, and Brainstem Regions in Alzheimer’s Disease. Curr Neurovasc Res. 2014:1–14.

12. Tang H, Mao X, Xie L, Greenberg DA, Jin K. Expression level of vascular endothelial growth factor in hippocampus is associated with cognitive impairment in patients with Alzheimer’s disease. Neurobiol Aging. 2013;34:1412–1415.

13. Thomas T, Miners S, Love S. Post-mortem assessment of hypoperfusion of cerebral cortex in Alzheimer’s disease and vascular dementia. Brain. 2015;138:1059–1069.

14. Garcia KO, Ornellas FLM, Martin PKM, Patti CL, Mello LE, Frussa-Filho R, et al. Therapeutic effects of the transplantation of VEGF overexpressing bone marrow mesenchymal stem cells in the hippocampus of murine model of Alzheimer’s disease. Front Aging Neurosci. 2014;6:6–30.

15. Hohman TJ, Bell SP, Jefferson AL. The Role of Vascular Endothelial Growth Factor in Neurodegeneration and Cognitive Decline: Exploring Interactions With Biomarkers of Alzheimer Disease. JAMA Neurol. 2015;72:520–529.

16. Mahoney ER, Dumitrescu L, Moore AM, Cambronero FE, De Jager PL, Koran MEI, et al. Brain expression of the vascular endothelial growth factor gene family in cognitive aging and Alzheimer’s disease. Mol Psychiatry. 2019. 22 July 2019. 10.1038/s41380-019-0458-5.

17. Libby JB, Seto M, Khan OA, Liu D, Petyuk VA, Oliver N, et al. Whole blood transcript and protein abundance of the vascular endothelial growth factor family relate to cognitive performance. Neurobiol Aging. 2023;124:11–17.

18. Seto M, Dumitrescu L, Mahoney ER, Sclafani AM, De Jager PL, Menon V, et al. Multi-omic characterization of brain changes in the vascular endothelial growth factor family during aging and Alzheimer’s disease. Neurobiol Aging. 2023;126:25–33.

19. Green GS, Fujita M, Yang H-S, Taga M, McCabe C, Cain A, et al. Cellular dynamics across aged human brains uncover a multicellular cascade leading to Alzheimer’s disease. 2023:2023.03.07.531493.

20. Fujita M, Gao Z, Zeng L, McCabe C, White CC, Ng B, et al. Cell subtype-specific effects of genetic variation in the Alzheimer’s disease brain. Nat Genet. 2024;56:605–614.

21. Bennett DA, Schneider JA, Arvanitakis Z, Wilson RS. Overview and findings from the religious orders study. Curr Alzheimer Res. 2012;9:628.

22. Bennett DA, Schneider JA, Buchman AS, Barnes LL, Boyle PA, Wilson RS. Overview and findings from the Rush Memory and Aging Project. Curr Alzheimer Res. 2012;9:646.

23. Bennett DA, Buchman AS, Boyle PA, Barnes LL, Wilson RS, Schneider JA. Religious Orders Study and Rush Memory and Aging Project. J Alzheimers Dis. 2018;64:S161–S189.

24. Fujita M, Gao Z, Zeng L, McCabe C, White CC, Ng B, et al. Cell-subtype specific effects of genetic variation in the aging and Alzheimer cortex. 2022.

25. Cain A, Taga M, McCabe C, Green GS, Hekselman I, White CC. Multicellular communities are perturbed in the aging human brain and Alzheimer’s disease. Nat Neurosci. 2023;26:1267–1280.

26. Habib N, Avraham-Davidi I, Basu A, Burks T, Shekhar K, Hofree M, et al. Massively parallel single-nucleus RNA-seq with DroNc-seq. Nat Methods. 2017;14:955–958.

27. Wilson RS, Boyle PA, Yu L, Barnes LL, Sytsma J, Buchman AS, et al. Temporal course and pathologic basis of unawareness of memory loss in dementia. Neurology. 2015;85:984–991.

28. He L, Davila-Velderrain J, Sumida TS, Hafler DA, Kellis M, Kulminski AM. NEBULA is a fast negative binomial mixed model for differential or co-expression analysis of large-scale multi-subject single-cell data. Commun Biol. 2021;4:629.

29. Jin S, Guerrero-Juarez CF, Zhang L, Chang I, Ramos R, Kuan C-H, et al. Inference and analysis of cell-cell communication using CellChat. Nat Commun. 2021;12:1088.

30. Hicklin DJ, Ellis LM. Role of the Vascular Endothelial Growth Factor Pathway in Tumor Growth and Angiogenesis. J Clin Oncol. 2005;23:1011–1027.

31. Shibuya M. Vascular Endothelial Growth Factor (VEGF) and Its Receptor (VEGFR) Signaling in Angiogenesis. Genes Cancer. 2011;2:1097–1105.

32. Jefferies WA, Price KA, Biron KE, Fenninger F, Pfeifer CG, Dickstein DL. Adjusting the compass: new insights into the role of angiogenesis in Alzheimer’s disease. Alzheimers Res Ther. 2013;5:64.

33. Lau S-F, Cao H, Fu AKY, Ip NY. Single-nucleus transcriptome analysis reveals dysregulation of angiogenic endothelial cells and neuroprotective glia in Alzheimer’s disease. Proc Natl Acad Sci. 2020;117:25800.

34. Tsartsalis S, Sleven H, Fancy N, Wessely F, Smith AM, Willumsen N, et al. A single nuclear transcriptomic characterisation of mechanisms responsible for impaired angiogenesis and blood-brain barrier function in Alzheimer’s disease. Nat Commun. 2024;15:2243.

35. Hunter M, Spiller KJ, Dominique MA, Xu H, Hunter FW, Fang TC, et al. Microglial transcriptome analysis in the rNLS8 mouse model of TDP-43 proteinopathy reveals discrete expression profiles associated with neurodegenerative progression and recovery. Acta Neuropathol Commun. 2021;9:140.

36. Stankovic I, Notaras M, Wolujewicz P, Lu T, Lis R, Ross ME, et al. Schizophrenia endothelial cells exhibit higher permeability and altered angiogenesis patterns in patient-derived organoids. Transl Psychiatry. 2024;14:1–15.

37. Desai BS, Schneider JA, Li J-L, Carvey PM, Hendey B. Evidence of angiogenic vessels in Alzheimer’s disease. J Neural Transm Vienna Austria 1996. 2009;116:587–597.

38. Vogel C, Bauer A, Wiesnet M, Preissner KT, Schaper W, Marti HH, et al. Flt-1, but not Flk-1 mediates hyperpermeability through activation of the PI3-K/Akt pathway. J Cell Physiol. 2007;212:236–243.

39. Gao L, Pan X, Zhang JH, Xia Y. Glial cells: an important switch for the vascular function of the central nervous system. Front Cell Neurosci. 2023;17.

40. Sun Q, Xu X, Wang T, Xu Z, Lu X, Li X, et al. Neurovascular Units and Neural-Glia Networks in Intracerebral Hemorrhage: from Mechanisms to Translation. Transl Stroke Res. 2021;12:447–460.

41. Daneman R, Prat A. The Blood–Brain Barrier. Cold Spring Harb Perspect Biol. 2015;7:a020412.

42. Wu D, Chen Q, Chen X, Han F, Chen Z, Wang Y. The blood–brain barrier: structure, regulation, and drug delivery. Signal Transduct Target Ther. 2023;8:1–27.

43. Zhang S, Kim B, Zhu X, Gui X, Wang Y, Lan Z, et al. Glial type specific regulation of CNS angiogenesis by HIFα-activated different signaling pathways. Nat Commun. 2020;11:2027.

44. Egensperger R, Kosel S, von Eitzen U, Graeber MB. Microglial activation in Alzheimer disease: Association with APOE genotype. Brain Pathol. 1998;8:439–447.

45. Maezawa I, Zimin PI, Wulff H, Jin LW. Amyloid-beta protein oligomer at low nanomolar concentrations activates microglia and induces microglial neurotoxicity. J Biol Chem;286:3693–3706.

46. Malik M, Parikh I, Vasquez JB, Smith C, Tai L, Bu G, et al. Genetics ignite focus on microglial inflammation in Alzheimer’s disease. Mol Neurodegener. 2015;10:52.

47. Nordengen K, Kirsebom BE, Henjum K, Selnes P, Gisladottir B, Wettergreen M, et al. Glial activation and inflammation along the Alzheimer’s disease continuum. J Neuroinflammation. 2019;16:46.

48. Rogers J, Strohmeyer R, Kovelowski CJ, Li R. Microglia and inflammatory mechanisms in the clearance of amyloid-beta peptide. Glia. 2002;40:260–269.

49. Ryu JK, Cho T, Choi HB, Wang YT, McLarnon JG. Microglial VEGF receptor response is an integral chemotactic component in Alzheimer’s disease pathology. J Neurosci. 2009;29:3–13.

50. Xiong F, Ge W, Ma C. Quantitative proteomics reveals distinct composition of amyloid plaques in Alzheimer’s disease. Alzheimers Dement. 2019;15:429–440.

51. Lau S-F, Wu W, Wong HY, Ouyang L, Qiao Y, Xu J, et al. The VCAM1–ApoE pathway directs microglial chemotaxis and alleviates Alzheimer’s disease pathology. Nat Aging. 2023:1–18.

52. Sawano A, Iwai S, Sakurai Y, Ito M, Shitara K, Nakahata T, et al. Flt-1, vascular endothelial growth factor receptor 1, is a novel cell surface marker for the lineage of monocyte-macrophages in humans. Blood. 2001;97:785–791.

53. Hiratsuka S, Minowa O, Kuno J, Noda T, Shibuya M. Flt-1 lacking the tyrosine kinase domain is sufficient for normal development and angiogenesis in mice. Proc Natl Acad Sci. 1998;95:9349–9354.

54. Kalafatakis I, Karagogeos D. Oligodendrocytes and Microglia: Key Players in Myelin Development, Damage and Repair. Biomolecules. 2021;11:1058.

55. Li Y, Zhang F, Nagai N, Tang Z, Zhang S, Scotney P, et al. VEGF-B inhibits apoptosis via VEGFR-1–mediated suppression of the expression of BH3-only protein genes in mice and rats. J Clin Invest. 2008;118:913.

56. Falk T, Yue X, Zhang S, McCourt AD, Yee BJ, Gonzalez RT, et al. Vascular endothelial growth factor-B is neuroprotective in an in vivo rat model of Parkinson’s disease. Neurosci Lett. 2011;496:43–47.

57. Caballero B, Sherman SJ, Falk T. Insights into the Mechanisms Involved in Protective Effects of VEGF-B in Dopaminergic Neurons. Park Dis. 2017;2017:4263795.

58. Guaiquil VH, Pan Z, Karagianni N, Fukuoka S, Alegre G, Rosenblatt MI. VEGF-B selectively regenerates injured peripheral neurons and restores sensory and trophic functions. Proc Natl Acad Sci U S A. 2014;111:17272–17277.

59. Fu H, Possenti A, Freer R, Nakano Y, Hernandez Villegas NC, Tang M, et al. A tau homeostasis signature is linked with the cellular and regional vulnerability of excitatory neurons to tau pathology. Nat Neurosci. 2019;22:47–56.

60. Kiss E, Gorgas K, Schlicksupp A, Groß D, Kins S, Kirsch J, et al. Biphasic Alteration of the Inhibitory Synapse Scaffold Protein Gephyrin in Early and Late Stages of an Alzheimer Disease Model. Am J Pathol. 2016;186:2279–2291.

61. Da Mesquita S, Papadopoulos Z, Dykstra T, Brase L, Farias FG, Wall M, et al. Meningeal lymphatics affect microglia responses and anti-Aβ immunotherapy. Nature. 2021;593:255–260.

62. Weijts BGMW. Atypical E2fs Control Lymphangiogenesis through Transcriptional Regulation of Ccbe1 and Flt4. PloS One. 2013;8:e73693.

