## Supplemental Table 1 for "Association of 10 VEGF Family Genes with Alzheimer’s Disease Endophenotypes at Single Cell Resolution"

**Supplemental Table 1. Association of 10 VEGF Gene Expression and Various AD Outcomes**

| Gene | Outcome | Cell Type | logFC | SE | P Value | FDR |
|---|---|---|---|---|---|---|
| *FLT1* | Diagnosis | Astrocytes | -0.022 | 0.081 | 0.783 | 0.926 |
| *FLT1* | Diagnosis | Endothelial cells | 0.195 | 0.055 | 0 | 0.014 |
| *FLT1* | Diagnosis | CUX2- Excitatory neurons | 0.015 | 0.083 | 0.86 | 0.944 |
| *FLT1* | Diagnosis | CUX2+ Excitatory neurons | -0.011 | 0.087 | 0.896 | 0.963 |
| *FLT1* | Diagnosis | Inhibitory neurons | -0.08 | 0.038 | 0.035 | 0.183 |
| *FLT1* | Diagnosis | Microglia | 0.452 | 0.113 | 0 | 0.009 |
| *FLT1* | Diagnosis | Oligodendrocytes | -0.054 | 0.085 | 0.528 | 0.773 |
| *FLT1* | Diagnosis | Oligodendrocyte precursor cells | 0.001 | 0.119 | 0.994 | 0.994 |
| *FLT4* | Diagnosis | Endothelial cells | 0.165 | 0.088 | 0.06 | 0.246 |
| *KDR* | Diagnosis | Endothelial cells | 0.094 | 0.08 | 0.24 | 0.552 |
| *KDR* | Diagnosis | Inhibitory neurons | 0.165 | 0.072 | 0.022 | 0.161 |
| *KDR* | Diagnosis | Oligodendrocytes | -0.009 | 0.059 | 0.882 | 0.958 |
| *NRP1* | Diagnosis | Astrocytes | -0.118 | 0.064 | 0.067 | 0.272 |
| *NRP1* | Diagnosis | Endothelial cells | 0.075 | 0.065 | 0.247 | 0.552 |
| *NRP1* | Diagnosis | CUX2- Excitatory neurons | -0.071 | 0.055 | 0.195 | 0.523 |
| *NRP1* | Diagnosis | CUX2+ Excitatory neurons | 0.059 | 0.034 | 0.084 | 0.316 |
| *NRP1* | Diagnosis | Inhibitory neurons | -0.007 | 0.033 | 0.833 | 0.937 |
| *NRP1* | Diagnosis | Microglia | 0.084 | 0.05 | 0.092 | 0.329 |
| *NRP1* | Diagnosis | Oligodendrocytes | -0.041 | 0.063 | 0.518 | 0.773 |
| *NRP1* | Diagnosis | Oligodendrocyte precursor cells | -0.014 | 0.066 | 0.827 | 0.937 |
| *NRP2* | Diagnosis | Astrocytes | -0.115 | 0.095 | 0.228 | 0.55 |
| *NRP2* | Diagnosis | Endothelial cells | 0.02 | 0.043 | 0.634 | 0.832 |
| *NRP2* | Diagnosis | CUX2- Excitatory neurons | -0.042 | 0.034 | 0.218 | 0.55 |
| *NRP2* | Diagnosis | CUX2+ Excitatory neurons | 0.025 | 0.041 | 0.533 | 0.773 |
| *NRP2* | Diagnosis | Inhibitory neurons | -0.069 | 0.032 | 0.034 | 0.181 |
| *NRP2* | Diagnosis | Microglia | -0.101 | 0.038 | 0.007 | 0.086 |
| *NRP2* | Diagnosis | Oligodendrocytes | -0.073 | 0.056 | 0.187 | 0.522 |
| *NRP2* | Diagnosis | Oligodendrocyte precursor cells | -0.087 | 0.075 | 0.247 | 0.552 |
| *PGF* | Diagnosis | Astrocytes | -0.09 | 0.105 | 0.388 | 0.647 |
| *PGF* | Diagnosis | Endothelial cells | 0.086 | 0.079 | 0.279 | 0.594 |
| *VEGFA* | Diagnosis | Astrocytes | 0.046 | 0.079 | 0.561 | 0.782 |
| *VEGFA* | Diagnosis | Endothelial cells | 0.207 | 0.096 | 0.032 | 0.181 |

| | | | | | | |
|---|---|---|---|---|---|---|
| *VEGFA* | Diagnosis | CUX2- Excitatory neurons | -0.047 | 0.055 | 0.391 | 0.647 |
| *VEGFA* | Diagnosis | CUX2+ Excitatory neurons | 0.057 | 0.059 | 0.331 | 0.634 |
| *VEGFA* | Diagnosis | Inhibitory neurons | -0.031 | 0.041 | 0.447 | 0.713 |
| *VEGFA* | Diagnosis | Microglia | 0.002 | 0.085 | 0.984 | 0.994 |
| *VEGFA* | Diagnosis | Oligodendrocytes | 0.079 | 0.082 | 0.334 | 0.634 |
| *VEGFA* | Diagnosis | Oligodendrocyte precursor cells | -0.024 | 0.098 | 0.805 | 0.926 |
| *VEGFB* | Diagnosis | Astrocytes | -0.004 | 0.032 | 0.908 | 0.963 |
| *VEGFB* | Diagnosis | Endothelial cells | 0.091 | 0.064 | 0.156 | 0.467 |
| *VEGFB* | Diagnosis | CUX2- Excitatory neurons | -0.048 | 0.046 | 0.29 | 0.604 |
| *VEGFB* | Diagnosis | CUX2+ Excitatory neurons | -0.013 | 0.051 | 0.801 | 0.926 |
| *VEGFB* | Diagnosis | Inhibitory neurons | -0.068 | 0.038 | 0.074 | 0.292 |
| *VEGFB* | Diagnosis | Microglia | -0.071 | 0.041 | 0.086 | 0.316 |
| *VEGFB* | Diagnosis | Oligodendrocytes | 0.082 | 0.039 | 0.039 | 0.192 |
| *VEGFB* | Diagnosis | Oligodendrocyte precursor cells | 0.003 | 0.033 | 0.925 | 0.975 |
| *VEGFC* | Diagnosis | Astrocytes | 0.072 | 0.055 | 0.193 | 0.522 |
| *VEGFC* | Diagnosis | Endothelial cells | 0.048 | 0.095 | 0.61 | 0.819 |
| *VEGFC* | Diagnosis | CUX2- Excitatory neurons | 0.18 | 0.066 | 0.007 | 0.086 |
| *VEGFC* | Diagnosis | Inhibitory neurons | 0.017 | 0.061 | 0.777 | 0.926 |
| *VEGFD* | Diagnosis | Astrocytes | -0.041 | 0.054 | 0.447 | 0.713 |
| *VEGFD* | Diagnosis | Endothelial cells | -0.197 | 0.127 | 0.12 | 0.4 |
| *VEGFD* | Diagnosis | CUX2- Excitatory neurons | 0.027 | 0.022 | 0.224 | 0.55 |
| *VEGFD* | Diagnosis | CUX2+ Excitatory neurons | 0.061 | 0.028 | 0.029 | 0.181 |
| *VEGFD* | Diagnosis | Inhibitory neurons | 0.031 | 0.027 | 0.256 | 0.562 |
| *VEGFD* | Diagnosis | Microglia | -0.019 | 0.109 | 0.862 | 0.944 |
| *VEGFD* | Diagnosis | Oligodendrocytes | 0.031 | 0.048 | 0.527 | 0.773 |
| *VEGFD* | Diagnosis | Oligodendrocyte precursor cells | 0.095 | 0.082 | 0.247 | 0.552 |
| *FLT1* | β-amyloid load | Astrocytes | 0.016 | 0.007 | 0.023 | 0.161 |
| *FLT1* | β-amyloid load | Endothelial cells | 0.009 | 0.005 | 0.087 | 0.316 |
| *FLT1* | β-amyloid load | CUX2- Excitatory neurons | -0.008 | 0.007 | 0.3 | 0.605 |
| *FLT1* | β-amyloid load | CUX2+ Excitatory neurons | 0.021 | 0.008 | 0.013 | 0.123 |
| *FLT1* | β-amyloid load | Inhibitory neurons | -0.001 | 0.004 | 0.799 | 0.926 |
| *FLT1* | β-amyloid load | Microglia | 0.053 | 0.011 | 0 | 0.001 |
| *FLT1* | β-amyloid load | Oligodendrocytes | 0.017 | 0.008 | 0.048 | 0.212 |
| *FLT1* | β-amyloid load | Oligodendrocyte precursor cells | 0.027 | 0.011 | 0.015 | 0.131 |
| *FLT4* | β-amyloid load | Endothelial cells | 0.027 | 0.008 | 0 | 0.013 |

| | | | | | | |
|---|---|---|---|---|---|---|
| *KDR* | β-amyloid load | Endothelial cells | -0.009 | 0.007 | 0.253 | 0.559 |
| *KDR* | β-amyloid load | Inhibitory neurons | -0.004 | 0.007 | 0.523 | 0.773 |
| *KDR* | β-amyloid load | Oligodendrocytes | 0.003 | 0.005 | 0.597 | 0.81 |
| *NRP1* | β-amyloid load | Astrocytes | -0.012 | 0.006 | 0.056 | 0.238 |
| *NRP1* | β-amyloid load | Endothelial cells | 0.013 | 0.006 | 0.026 | 0.177 |
| *NRP1* | β-amyloid load | CUX2- Excitatory neurons | -0.011 | 0.005 | 0.023 | 0.161 |
| *NRP1* | β-amyloid load | CUX2+ Excitatory neurons | 0.006 | 0.003 | 0.045 | 0.202 |
| *NRP1* | β-amyloid load | Inhibitory neurons | 0.004 | 0.003 | 0.233 | 0.55 |
| *NRP1* | β-amyloid load | Microglia | 0.008 | 0.005 | 0.075 | 0.292 |
| *NRP1* | β-amyloid load | Oligodendrocytes | 0.014 | 0.007 | 0.029 | 0.181 |
| *NRP1* | β-amyloid load | Oligodendrocyte precursor cells | 0.018 | 0.006 | 0.002 | 0.046 |
| *NRP2* | β-amyloid load | Astrocytes | 0 | 0.01 | 0.967 | 0.988 |
| *NRP2* | β-amyloid load | Endothelial cells | 0.008 | 0.004 | 0.051 | 0.224 |
| *NRP2* | β-amyloid load | CUX2- Excitatory neurons | 0 | 0.003 | 0.897 | 0.963 |
| *NRP2* | β-amyloid load | CUX2+ Excitatory neurons | 0.005 | 0.004 | 0.188 | 0.522 |
| *NRP2* | β-amyloid load | Inhibitory neurons | 0 | 0.003 | 0.88 | 0.958 |
| *NRP2* | β-amyloid load | Microglia | 0.007 | 0.004 | 0.035 | 0.183 |
| *NRP2* | β-amyloid load | Oligodendrocytes | 0.005 | 0.005 | 0.379 | 0.647 |
| *NRP2* | β-amyloid load | Oligodendrocyte precursor cells | 0.006 | 0.007 | 0.383 | 0.647 |
| *PGF* | β-amyloid load | Astrocytes | 0.007 | 0.01 | 0.5 | 0.768 |
| *PGF* | β-amyloid load | Endothelial cells | 0.018 | 0.007 | 0.015 | 0.131 |
| *VEGFA* | β-amyloid load | Astrocytes | -0.003 | 0.007 | 0.687 | 0.882 |
| *VEGFA* | β-amyloid load | Endothelial cells | -0.001 | 0.009 | 0.908 | 0.963 |
| *VEGFA* | β-amyloid load | CUX2- Excitatory neurons | -0.01 | 0.005 | 0.056 | 0.238 |
| *VEGFA* | β-amyloid load | CUX2+ Excitatory neurons | -0.006 | 0.006 | 0.289 | 0.604 |
| *VEGFA* | β-amyloid load | Inhibitory neurons | -0.012 | 0.004 | 0.003 | 0.056 |
| *VEGFA* | β-amyloid load | Microglia | 0.006 | 0.008 | 0.503 | 0.768 |
| *VEGFA* | β-amyloid load | Oligodendrocytes | -0.001 | 0.008 | 0.935 | 0.977 |
| *VEGFA* | β-amyloid load | Oligodendrocyte precursor cells | -0.019 | 0.009 | 0.04 | 0.192 |
| *VEGFB* | β-amyloid load | Astrocytes | 0.005 | 0.003 | 0.144 | 0.443 |
| *VEGFB* | β-amyloid load | Endothelial cells | 0.002 | 0.006 | 0.772 | 0.925 |
| *VEGFB* | β-amyloid load | CUX2- Excitatory neurons | -0.005 | 0.004 | 0.217 | 0.55 |
| *VEGFB* | β-amyloid load | CUX2+ Excitatory neurons | 0 | 0.005 | 0.936 | 0.977 |
| *VEGFB* | β-amyloid load | Inhibitory neurons | -0.012 | 0.004 | 0.001 | 0.019 |
| *VEGFB* | β-amyloid load | Microglia | 0.009 | 0.004 | 0.026 | 0.177 |

| | | | | | | |
|---|---|---|---|---|---|---|
| *VEGFB* | β-amyloid load | Oligodendrocytes | 0.014 | 0.004 | 0 | 0.009 |
| *VEGFB* | β-amyloid load | Oligodendrocyte precursor cells | 0.007 | 0.003 | 0.029 | 0.181 |
| *VEGFC* | β-amyloid load | Astrocytes | 0.013 | 0.005 | 0.008 | 0.091 |
| *VEGFC* | β-amyloid load | Endothelial cells | -0.001 | 0.009 | 0.946 | 0.98 |
| *VEGFC* | β-amyloid load | CUX2- Excitatory neurons | -0.004 | 0.006 | 0.571 | 0.789 |
| *VEGFC* | β-amyloid load | Inhibitory neurons | -0.01 | 0.006 | 0.101 | 0.352 |
| *VEGFD* | β-amyloid load | Astrocytes | 0.016 | 0.005 | 0.001 | 0.029 |
| *VEGFD* | β-amyloid load | Endothelial cells | 0.015 | 0.011 | 0.181 | 0.522 |
| *VEGFD* | β-amyloid load | CUX2- Excitatory neurons | 0.001 | 0.002 | 0.674 | 0.872 |
| *VEGFD* | β-amyloid load | CUX2+ Excitatory neurons | 0.005 | 0.002 | 0.031 | 0.181 |
| *VEGFD* | β-amyloid load | Inhibitory neurons | 0.002 | 0.003 | 0.387 | 0.647 |
| *VEGFD* | β-amyloid load | Microglia | 0.015 | 0.01 | 0.114 | 0.384 |
| *VEGFD* | β-amyloid load | Oligodendrocytes | 0.01 | 0.004 | 0.033 | 0.181 |
| *VEGFD* | β-amyloid load | Oligodendrocyte precursor cells | 0.015 | 0.008 | 0.057 | 0.238 |
| *FLT1* | Cognition trajectory | Astrocytes | 0.455 | 0.343 | 0.184 | 0.522 |
| *FLT1* | Cognition trajectory | Endothelial cells | -0.771 | 0.248 | 0.002 | 0.041 |
| *FLT1* | Cognition trajectory | Inhibitory neurons | 0.44 | 0.164 | 0.007 | 0.086 |
| *FLT1* | Cognition trajectory | Microglia | -1.964 | 0.504 | 0 | 0.009 |
| *FLT1* | Cognition trajectory | Oligodendrocyte precursor cells | 0.666 | 0.527 | 0.207 | 0.539 |
| *FLT1* | Cognition trajectory | CUX2- Excitatory neurons | 0.167 | 0.343 | 0.626 | 0.829 |
| *FLT1* | Cognition trajectory | CUX2+ Excitatory neurons | -0.022 | 0.401 | 0.957 | 0.987 |
| *FLT1* | Cognition trajectory | Oligodendrocytes | 0.449 | 0.388 | 0.247 | 0.552 |
| *FLT4* | Cognition trajectory | Endothelial cells | -1.089 | 0.373 | 0.004 | 0.056 |
| *KDR* | Cognition trajectory | Endothelial cells | 0.093 | 0.346 | 0.787 | 0.926 |
| *KDR* | Cognition trajectory | Inhibitory neurons | -0.381 | 0.316 | 0.228 | 0.55 |
| *KDR* | Cognition trajectory | Oligodendrocytes | 0.044 | 0.256 | 0.862 | 0.944 |
| *NRP1* | Cognition trajectory | Astrocytes | 0.825 | 0.286 | 0.004 | 0.06 |
| *NRP1* | Cognition trajectory | Endothelial cells | -0.29 | 0.287 | 0.311 | 0.606 |
| *NRP1* | Cognition trajectory | Inhibitory neurons | 0.1 | 0.146 | 0.493 | 0.768 |
| *NRP1* | Cognition trajectory | Microglia | -0.269 | 0.213 | 0.208 | 0.539 |
| *NRP1* | Cognition trajectory | Oligodendrocyte precursor cells | 0.209 | 0.284 | 0.462 | 0.732 |
| *NRP1* | Cognition trajectory | CUX2- Excitatory neurons | 0.495 | 0.232 | 0.033 | 0.181 |
| *NRP1* | Cognition trajectory | CUX2+ Excitatory neurons | -0.15 | 0.146 | 0.306 | 0.606 |
| *NRP1* | Cognition trajectory | Oligodendrocytes | 0.004 | 0.304 | 0.99 | 0.994 |
| *NRP2* | Cognition trajectory | Astrocytes | 0.747 | 0.437 | 0.087 | 0.316 |

| | | | | | | |
|---|---|---|---|---|---|---|
| *NRP2* | Cognition trajectory | Endothelial cells | -0.169 | 0.195 | 0.384 | 0.647 |
| *NRP2* | Cognition trajectory | Inhibitory neurons | 0.289 | 0.143 | 0.043 | 0.202 |
| *NRP2* | Cognition trajectory | Microglia | 0.484 | 0.165 | 0.003 | 0.056 |
| *NRP2* | Cognition trajectory | Oligodendrocyte precursor cells | 0.347 | 0.33 | 0.294 | 0.605 |
| *NRP2* | Cognition trajectory | CUX2- Excitatory neurons | 0.095 | 0.156 | 0.545 | 0.775 |
| *NRP2* | Cognition trajectory | CUX2+ Excitatory neurons | -0.391 | 0.166 | 0.019 | 0.146 |
| *NRP2* | Cognition trajectory | Oligodendrocytes | 0.217 | 0.246 | 0.378 | 0.647 |
| *PGF* | Cognition trajectory | Astrocytes | 0.853 | 0.499 | 0.087 | 0.316 |
| *PGF* | Cognition trajectory | Endothelial cells | -0.327 | 0.35 | 0.35 | 0.639 |
| *VEGFA* | Cognition trajectory | Astrocytes | -0.223 | 0.337 | 0.508 | 0.772 |
| *VEGFA* | Cognition trajectory | Endothelial cells | -0.934 | 0.396 | 0.018 | 0.146 |
| *VEGFA* | Cognition trajectory | Inhibitory neurons | 0.175 | 0.183 | 0.339 | 0.637 |
| *VEGFA* | Cognition trajectory | Microglia | 0.256 | 0.391 | 0.512 | 0.773 |
| *VEGFA* | Cognition trajectory | Oligodendrocyte precursor cells | 0.623 | 0.413 | 0.131 | 0.423 |
| *VEGFA* | Cognition trajectory | CUX2- Excitatory neurons | 0.315 | 0.241 | 0.192 | 0.522 |
| *VEGFA* | Cognition trajectory | CUX2+ Excitatory neurons | -0.102 | 0.266 | 0.702 | 0.893 |
| *VEGFA* | Cognition trajectory | Oligodendrocytes | -0.531 | 0.387 | 0.17 | 0.498 |
| *VEGFB* | Cognition trajectory | Astrocytes | -0.163 | 0.146 | 0.266 | 0.579 |
| *VEGFB* | Cognition trajectory | Endothelial cells | -0.244 | 0.28 | 0.384 | 0.647 |
| *VEGFB* | Cognition trajectory | Inhibitory neurons | 0.266 | 0.174 | 0.126 | 0.411 |
| *VEGFB* | Cognition trajectory | Microglia | -0.046 | 0.187 | 0.804 | 0.926 |
| *VEGFB* | Cognition trajectory | Oligodendrocyte precursor cells | -0.098 | 0.155 | 0.527 | 0.773 |
| *VEGFB* | Cognition trajectory | CUX2- Excitatory neurons | 0.065 | 0.206 | 0.754 | 0.912 |
| *VEGFB* | Cognition trajectory | CUX2+ Excitatory neurons | -0.361 | 0.235 | 0.123 | 0.407 |
| *VEGFB* | Cognition trajectory | Oligodendrocytes | -0.561 | 0.166 | 0.001 | 0.019 |
| *VEGFC* | Cognition trajectory | Astrocytes | -0.169 | 0.234 | 0.469 | 0.74 |
| *VEGFC* | Cognition trajectory | Endothelial cells | 0.61 | 0.417 | 0.143 | 0.443 |
| *VEGFC* | Cognition trajectory | Inhibitory neurons | -0.185 | 0.273 | 0.499 | 0.768 |
| *VEGFC* | Cognition trajectory | CUX2- Excitatory neurons | -0.754 | 0.291 | 0.01 | 0.099 |
| *VEGFD* | Cognition trajectory | Astrocytes | 0.241 | 0.227 | 0.288 | 0.604 |
| *VEGFD* | Cognition trajectory | Endothelial cells | 0.619 | 0.57 | 0.277 | 0.594 |
| *VEGFD* | Cognition trajectory | Inhibitory neurons | -0.076 | 0.119 | 0.523 | 0.773 |
| *VEGFD* | Cognition trajectory | Microglia | 0.108 | 0.472 | 0.819 | 0.935 |
| *VEGFD* | Cognition trajectory | Oligodendrocyte precursor cells | -0.192 | 0.361 | 0.595 | 0.81 |
| *VEGFD* | Cognition trajectory | CUX2- Excitatory neurons | -0.08 | 0.099 | 0.42 | 0.684 |

| | | | | | | |
|---|---|---|---|---|---|---|
| *VEGFD* | Cognition trajectory | CUX2+ Excitatory neurons | -0.125 | 0.12 | 0.296 | 0.605 |
| *VEGFD* | Cognition trajectory | Oligodendrocytes | -0.041 | 0.208 | 0.844 | 0.941 |
| *FLT1* | Last cognition before death | Astrocytes | -0.004 | 0.033 | 0.91 | 0.963 |
| *FLT1* | Last cognition before death | Endothelial cells | -0.089 | 0.024 | 0 | 0.009 |
| *FLT1* | Last cognition before death | Inhibitory neurons | 0.034 | 0.016 | 0.031 | 0.181 |
| *FLT1* | Last cognition before death | Microglia | -0.176 | 0.046 | 0 | 0.009 |
| *FLT1* | Last cognition before death | Oligodendrocyte precursor cells | 0.073 | 0.052 | 0.158 | 0.468 |
| *FLT1* | Last cognition before death | CUX2- Excitatory neurons | -0.001 | 0.033 | 0.964 | 0.988 |
| *FLT1* | Last cognition before death | CUX2+ Excitatory neurons | -0.013 | 0.038 | 0.731 | 0.901 |
| *FLT1* | Last cognition before death | Oligodendrocytes | 0.004 | 0.038 | 0.906 | 0.963 |
| *FLT4* | Last cognition before death | Endothelial cells | -0.102 | 0.035 | 0.003 | 0.056 |
| *KDR* | Last cognition before death | Endothelial cells | -0.032 | 0.032 | 0.31 | 0.606 |
| *KDR* | Last cognition before death | Inhibitory neurons | -0.058 | 0.031 | 0.058 | 0.242 |
| *KDR* | Last cognition before death | Oligodendrocytes | 0.008 | 0.024 | 0.746 | 0.909 |
| *NRP1* | Last cognition before death | Astrocytes | 0.01 | 0.027 | 0.716 | 0.901 |
| *NRP1* | Last cognition before death | Endothelial cells | -0.025 | 0.028 | 0.357 | 0.645 |
| *NRP1* | Last cognition before death | Inhibitory neurons | 0.005 | 0.014 | 0.729 | 0.901 |
| *NRP1* | Last cognition before death | Microglia | -0.018 | 0.021 | 0.381 | 0.647 |
| *NRP1* | Last cognition before death | Oligodendrocyte precursor cells | 0.001 | 0.027 | 0.979 | 0.993 |
| *NRP1* | Last cognition before death | CUX2- Excitatory neurons | 0.045 | 0.022 | 0.045 | 0.202 |
| *NRP1* | Last cognition before death | CUX2+ Excitatory neurons | -0.009 | 0.014 | 0.532 | 0.773 |
| *NRP1* | Last cognition before death | Oligodendrocytes | -0.018 | 0.03 | 0.553 | 0.78 |
| *NRP2* | Last cognition before death | Astrocytes | -0.002 | 0.04 | 0.967 | 0.988 |
| *NRP2* | Last cognition before death | Endothelial cells | -0.024 | 0.019 | 0.19 | 0.522 |
| *NRP2* | Last cognition before death | Inhibitory neurons | 0.016 | 0.013 | 0.222 | 0.55 |
| *NRP2* | Last cognition before death | Microglia | 0.042 | 0.016 | 0.008 | 0.09 |
| *NRP2* | Last cognition before death | Oligodendrocyte precursor cells | 0.019 | 0.032 | 0.544 | 0.775 |
| *NRP2* | Last cognition before death | CUX2- Excitatory neurons | 0.001 | 0.015 | 0.943 | 0.98 |
| *NRP2* | Last cognition before death | CUX2+ Excitatory neurons | -0.035 | 0.016 | 0.028 | 0.181 |
| *NRP2* | Last cognition before death | Oligodendrocytes | -0.03 | 0.023 | 0.188 | 0.522 |
| *PGF* | Last cognition before death | Astrocytes | 0.027 | 0.045 | 0.545 | 0.775 |
| *PGF* | Last cognition before death | Endothelial cells | -0.03 | 0.034 | 0.377 | 0.647 |
| *VEGFA* | Last cognition before death | Astrocytes | -0.035 | 0.03 | 0.246 | 0.552 |
| *VEGFA* | Last cognition before death | Endothelial cells | -0.105 | 0.038 | 0.005 | 0.073 |
| *VEGFA* | Last cognition before death | Inhibitory neurons | 0.007 | 0.017 | 0.691 | 0.883 |

| | | | | | | |
|---|---|---|---|---|---|---|
| *VEGFA* | Last cognition before death | Microglia | -0.035 | 0.037 | 0.343 | 0.637 |
| *VEGFA* | Last cognition before death | Oligodendrocyte precursor cells | 0.022 | 0.036 | 0.542 | 0.775 |
| *VEGFA* | Last cognition before death | CUX2- Excitatory neurons | 0.004 | 0.022 | 0.859 | 0.944 |
| *VEGFA* | Last cognition before death | CUX2+ Excitatory neurons | -0.031 | 0.024 | 0.201 | 0.535 |
| *VEGFA* | Last cognition before death | Oligodendrocytes | -0.076 | 0.037 | 0.04 | 0.192 |
| *VEGFB* | Last cognition before death | Astrocytes | -0.01 | 0.014 | 0.484 | 0.759 |
| *VEGFB* | Last cognition before death | Endothelial cells | -0.044 | 0.027 | 0.107 | 0.365 |
| *VEGFB* | Last cognition before death | Inhibitory neurons | 0.02 | 0.016 | 0.228 | 0.55 |
| *VEGFB* | Last cognition before death | Microglia | -0.009 | 0.018 | 0.605 | 0.816 |
| *VEGFB* | Last cognition before death | Oligodendrocyte precursor cells | -0.009 | 0.015 | 0.554 | 0.78 |
| *VEGFB* | Last cognition before death | CUX2- Excitatory neurons | 0.007 | 0.02 | 0.738 | 0.904 |
| *VEGFB* | Last cognition before death | CUX2+ Excitatory neurons | -0.027 | 0.023 | 0.231 | 0.55 |
| *VEGFB* | Last cognition before death | Oligodendrocytes | -0.038 | 0.016 | 0.017 | 0.139 |
| *VEGFC* | Last cognition before death | Astrocytes | -0.027 | 0.023 | 0.241 | 0.552 |
| *VEGFC* | Last cognition before death | Endothelial cells | 0.008 | 0.038 | 0.832 | 0.937 |
| *VEGFC* | Last cognition before death | Inhibitory neurons | -0.023 | 0.026 | 0.368 | 0.647 |
| *VEGFC* | Last cognition before death | CUX2- Excitatory neurons | -0.074 | 0.028 | 0.007 | 0.086 |
| *VEGFD* | Last cognition before death | Astrocytes | 0.024 | 0.022 | 0.292 | 0.605 |
| *VEGFD* | Last cognition before death | Endothelial cells | 0.097 | 0.057 | 0.085 | 0.316 |
| *VEGFD* | Last cognition before death | Inhibitory neurons | -0.009 | 0.011 | 0.447 | 0.713 |
| *VEGFD* | Last cognition before death | Microglia | 0.019 | 0.045 | 0.67 | 0.872 |
| *VEGFD* | Last cognition before death | Oligodendrocyte precursor cells | -0.033 | 0.034 | 0.325 | 0.629 |
| *VEGFD* | Last cognition before death | CUX2- Excitatory neurons | -0.01 | 0.01 | 0.307 | 0.606 |
| *VEGFD* | Last cognition before death | CUX2+ Excitatory neurons | -0.01 | 0.011 | 0.393 | 0.647 |
| *VEGFD* | Last cognition before death | Oligodendrocytes | -0.006 | 0.019 | 0.763 | 0.918 |
| *FLT1* | Tau density | Astrocytes | 0 | 0.005 | 0.979 | 0.993 |
| *FLT1* | Tau density | CUX2- Excitatory neurons | -0.004 | 0.004 | 0.347 | 0.637 |
| *FLT1* | Tau density | CUX2+ Excitatory neurons | 0.008 | 0.005 | 0.153 | 0.463 |
| *FLT1* | Tau density | Endothelial cells | 0.003 | 0.003 | 0.384 | 0.647 |
| *FLT1* | Tau density | Inhibitory neurons | -0.003 | 0.002 | 0.233 | 0.55 |
| *FLT1* | Tau density | Microglia | 0.02 | 0.007 | 0.006 | 0.077 |
| *FLT1* | Tau density | Oligodendrocytes | 0.001 | 0.005 | 0.797 | 0.926 |
| *FLT1* | Tau density | Oligodendrocyte precursor cells | 0.002 | 0.007 | 0.725 | 0.901 |
| *FLT4* | Tau density | Endothelial cells | 0.011 | 0.005 | 0.016 | 0.134 |
| *KDR* | Tau density | Endothelial cells | -0.001 | 0.005 | 0.818 | 0.935 |

| | | | | | | |
|---|---|---|---|---|---|---|
| *KDR* | Tau density | Inhibitory neurons | 0.003 | 0.004 | 0.496 | 0.768 |
| *KDR* | Tau density | Oligodendrocytes | 0 | 0.003 | 0.935 | 0.977 |
| *NRP1* | Tau density | Astrocytes | -0.014 | 0.004 | 0 | 0.009 |
| *NRP1* | Tau density | CUX2- Excitatory neurons | -0.003 | 0.003 | 0.358 | 0.645 |
| *NRP1* | Tau density | CUX2+ Excitatory neurons | 0.001 | 0.002 | 0.733 | 0.901 |
| *NRP1* | Tau density | Endothelial cells | 0.004 | 0.004 | 0.345 | 0.637 |
| *NRP1* | Tau density | Inhibitory neurons | 0.001 | 0.002 | 0.568 | 0.788 |
| *NRP1* | Tau density | Microglia | 0.002 | 0.003 | 0.561 | 0.782 |
| *NRP1* | Tau density | Oligodendrocytes | 0.001 | 0.004 | 0.837 | 0.937 |
| *NRP1* | Tau density | Oligodendrocyte precursor cells | -0.001 | 0.004 | 0.847 | 0.942 |
| *NRP2* | Tau density | Astrocytes | -0.006 | 0.006 | 0.27 | 0.583 |
| *NRP2* | Tau density | CUX2- Excitatory neurons | -0.001 | 0.002 | 0.797 | 0.926 |
| *NRP2* | Tau density | CUX2+ Excitatory neurons | 0.005 | 0.002 | 0.039 | 0.192 |
| *NRP2* | Tau density | Endothelial cells | 0.001 | 0.003 | 0.79 | 0.926 |
| *NRP2* | Tau density | Inhibitory neurons | -0.004 | 0.002 | 0.019 | 0.147 |
| *NRP2* | Tau density | Microglia | -0.004 | 0.002 | 0.105 | 0.362 |
| *NRP2* | Tau density | Oligodendrocytes | -0.002 | 0.003 | 0.647 | 0.845 |
| *NRP2* | Tau density | Oligodendrocyte precursor cells | -0.002 | 0.004 | 0.729 | 0.901 |
| *PGF* | Tau density | Astrocytes | -0.015 | 0.007 | 0.04 | 0.192 |
| *PGF* | Tau density | Endothelial cells | 0.007 | 0.005 | 0.137 | 0.435 |
| *VEGFA* | Tau density | Astrocytes | -0.001 | 0.005 | 0.894 | 0.963 |
| *VEGFA* | Tau density | CUX2- Excitatory neurons | -0.002 | 0.003 | 0.623 | 0.829 |
| *VEGFA* | Tau density | CUX2+ Excitatory neurons | 0.002 | 0.004 | 0.576 | 0.792 |
| *VEGFA* | Tau density | Endothelial cells | 0.008 | 0.006 | 0.147 | 0.448 |
| *VEGFA* | Tau density | Inhibitory neurons | 0.001 | 0.002 | 0.732 | 0.901 |
| *VEGFA* | Tau density | Microglia | 0.003 | 0.005 | 0.591 | 0.808 |
| *VEGFA* | Tau density | Oligodendrocytes | 0.005 | 0.005 | 0.306 | 0.606 |
| *VEGFA* | Tau density | Oligodendrocyte precursor cells | 0.003 | 0.006 | 0.617 | 0.825 |
| *VEGFB* | Tau density | Astrocytes | -0.001 | 0.002 | 0.755 | 0.912 |
| *VEGFB* | Tau density | CUX2- Excitatory neurons | -0.001 | 0.003 | 0.837 | 0.937 |
| *VEGFB* | Tau density | CUX2+ Excitatory neurons | 0.005 | 0.003 | 0.095 | 0.337 |
| *VEGFB* | Tau density | Endothelial cells | 0.004 | 0.004 | 0.334 | 0.634 |
| *VEGFB* | Tau density | Inhibitory neurons | -0.003 | 0.002 | 0.142 | 0.443 |
| *VEGFB* | Tau density | Microglia | 0 | 0.003 | 0.992 | 0.994 |
| *VEGFB* | Tau density | Oligodendrocytes | 0.006 | 0.002 | 0.013 | 0.123 |

| | | | | | | |
|---|---|---|---|---|---|---|
| *VEGFB* | Tau density | Oligodendrocyte precursor cells | -0.001 | 0.002 | 0.629 | 0.829 |
| *VEGFC* | Tau density | Astrocytes | 0.004 | 0.003 | 0.204 | 0.538 |
| *VEGFC* | Tau density | CUX2- Excitatory neurons | 0.009 | 0.004 | 0.013 | 0.123 |
| *VEGFC* | Tau density | Endothelial cells | -0.012 | 0.006 | 0.034 | 0.181 |
| *VEGFC* | Tau density | Inhibitory neurons | 0.001 | 0.004 | 0.687 | 0.882 |
| *VEGFD* | Tau density | Astrocytes | 0.003 | 0.003 | 0.382 | 0.647 |
| *VEGFD* | Tau density | CUX2- Excitatory neurons | 0 | 0.001 | 0.71 | 0.9 |
| *VEGFD* | Tau density | CUX2+ Excitatory neurons | 0.001 | 0.002 | 0.342 | 0.637 |
| *VEGFD* | Tau density | Endothelial cells | -0.006 | 0.008 | 0.432 | 0.7 |
| *VEGFD* | Tau density | Inhibitory neurons | 0.001 | 0.002 | 0.368 | 0.647 |
| *VEGFD* | Tau density | Microglia | -0.005 | 0.006 | 0.413 | 0.677 |
| *VEGFD* | Tau density | Oligodendrocytes | 0.003 | 0.003 | 0.231 | 0.55 |
| *VEGFD* | Tau density | Oligodendrocyte precursor cells | 0.005 | 0.005 | 0.3 | 0.605 |

*logFC values of diagnosis is log2 fold change of gene expression comparing AD to cognitive normal participants.
