## Supplemental Table 2 for "Association of 10 VEGF Family Genes with Alzheimer’s Disease Endophenotypes at Single Cell Resolution"

**Supplemental Table 2. VEGF Signaling Pathway Invesgated In This Study**

| Ligand | Receptor | Curation Source |
|---|---|---|
| VEGFA | FLT1 | CellChatDB |
| VEGFA | KDR | CellChatDB |
| VEGFB | FLT1 | CellChatDB |
| VEGFC | FLT4 | CellChatDB |
| VEGFC | KDR | CellChatDB |
| VEGFD | FLT4 | CellChatDB |
| VEGFD | KDR | CellChatDB |
| PGF | FLT1 | CellChatDB |
| VEGFA | FLT1_KDR | CellChatDB |
| VEGFC | FLT4_KDR | CellChatDB |
| VEGFD | FLT4_KDR | CellChatDB |
| VEGFD | ITGA4 | CellTalkDB |
| VEGFD | ITGB1 | CellTalkDB |
| VEGFD | NRP2 | CellTalkDB |
| VEGFB | RET | CellTalkDB |
| VEGFC | LYVE1 | CellTalkDB |
| VEGFA | ITGAV | CellTalkDB |
| VEGFC | FLT4 | CellTalkDB |
| VEGFD | FLT4 | CellTalkDB |
| VEGFA | TYRO3 | CellTalkDB |
| VEGFC | KDR | CellTalkDB |
| VEGFD | KDR | CellTalkDB |
| VEGFA | KDR | CellTalkDB |
| VEGFA | GPC1 | CellTalkDB |
| VEGFC | ITGA9 | CellTalkDB |
| VEGFD | ITGA9 | CellTalkDB |
| VEGFA | ITGA9 | CellTalkDB |
| VEGFB | NRP1 | CellTalkDB |
| PGF | NRP1 | CellTalkDB |
| VEGFA | NRP1 | CellTalkDB |
| VEGFA | EGFR | CellTalkDB |
| VEGFC | FLT1 | CellTalkDB |
| VEGFD | FLT1 | CellTalkDB |
| VEGFD | ITGA1 | CellTalkDB |
| VEGFD | ITGA5 | CellTalkDB |
| VEGFD | ITGA2 | CellTalkDB |
| VEGFA | RET | CellTalkDB |
| VEGFC | NRP2 | CellTalkDB |
| VEGFA | NRP2 | CellTalkDB |
| PGF | NRP2 | CellTalkDB |
| VEGFA | EPHB2 | CellTalkDB |
| VEGFC | ITGB1 | CellTalkDB |
| VEGFA | ITGB1 | CellTalkDB |
| VEGFA | SIRPA | CellTalkDB |
| VEGFC | CCBE1 | CellTalkDB |
| VEGFA | ITGB3 | CellTalkDB |
| VEGFA | GRIN2B | CellTalkDB |
