## Supplemental Table 3 for "Association of 10 VEGF Family Genes with Alzheimer’s Disease Endophenotypes at Single Cell Resolution"

**Supplemental Table 3. Communication Strength Comparison of Enriched VEGFA-Receptor Pairs Among 10 VEGFs Between AD And Cognition Normal Cognition Groups**

| Source | Target | Ligand | Receptor | Original Probability | Adjusted Probability* | Group |
|---|---|---|---|---|---|---|
| ast | endo | VEGFA | FLT1 | 1.89E-06 | 0.07587518 | Normal Cognition |
| ast | ast | VEGFA | ITGAV | 4.72E-07 | 0.06865381 | Normal Cognition |
| ast | endo | VEGFA | ITGAV | 1.18E-07 | 0.06268756 | Normal Cognition |
| ast | excit | VEGFA | ITGAV | 1.18E-07 | 0.06268756 | Normal Cognition |
| ast | inhib | VEGFA | ITGAV | 1.18E-07 | 0.06268756 | Normal Cognition |
| ast | mic | VEGFA | ITGAV | 1.18E-07 | 0.06268756 | Normal Cognition |
| ast | opc | VEGFA | ITGAV | 4.72E-07 | 0.06865381 | Normal Cognition |
| ast | excit | VEGFA | TYRO3 | 4.72E-07 | 0.06865381 | Normal Cognition |
| ast | oligo | VEGFA | TYRO3 | 1.18E-07 | 0.06268756 | Normal Cognition |
| ast | opc | VEGFA | TYRO3 | 1.18E-07 | 0.06268756 | Normal Cognition |
| ast | ast | VEGFA | GPC1 | 1.18E-07 | 0.06268756 | Normal Cognition |
| ast | excit | VEGFA | GPC1 | 3.54E-07 | 0.06732413 | Normal Cognition |
| ast | inhib | VEGFA | GPC1 | 1.18E-07 | 0.06268756 | Normal Cognition |
| ast | opc | VEGFA | GPC1 | 1.18E-07 | 0.06268756 | Normal Cognition |
| ast | ast | VEGFA | ITGA9 | 1.18E-07 | 0.06268756 | Normal Cognition |
| ast | excit | VEGFA | ITGA9 | 4.72E-07 | 0.06865381 | Normal Cognition |
| ast | inhib | VEGFA | ITGA9 | 2.36E-07 | 0.06553518 | Normal Cognition |
| ast | mic | VEGFA | ITGA9 | 1.18E-07 | 0.06268756 | Normal Cognition |
| ast | opc | VEGFA | ITGA9 | 1.18E-07 | 0.06268756 | Normal Cognition |
| ast | ast | VEGFA | NRP1 | 2.36E-07 | 0.06553518 | Normal Cognition |
| ast | endo | VEGFA | NRP1 | 1.18E-07 | 0.06268756 | Normal Cognition |
| ast | excit | VEGFA | NRP1 | 1.18E-07 | 0.06268756 | Normal Cognition |
| ast | mic | VEGFA | NRP1 | 1.18E-07 | 0.06268756 | Normal Cognition |
| ast | ast | VEGFA | EGFR | 5.90E-07 | 0.06972193 | Normal Cognition |
| ast | endo | VEGFA | EGFR | 4.72E-07 | 0.06865381 | Normal Cognition |
| ast | inhib | VEGFA | EGFR | 3.54E-07 | 0.06732413 | Normal Cognition |
| ast | opc | VEGFA | EGFR | 4.72E-07 | 0.06865381 | Normal Cognition |
| ast | endo | VEGFA | FLT1 | 1.89E-06 | 0.07587518 | Normal Cognition |
| ast | endo | VEGFA | NRP2 | 1.18E-07 | 0.06268756 | Normal Cognition |
| ast | excit | VEGFA | EPHB2 | 1.18E-07 | 0.06268756 | Normal Cognition |
| ast | inhib | VEGFA | EPHB2 | 1.18E-07 | 0.06268756 | Normal Cognition |

| | | | | | | |
|---|---|---|---|---|---|---|
| ast | ast | VEGFA | ITGB1 | 1.18E-07 | 0.06268756 | Normal Cognition |
| ast | endo | VEGFA | ITGB1 | 4.72E-07 | 0.06865381 | Normal Cognition |
| ast | excit | VEGFA | ITGB1 | 1.18E-07 | 0.06268756 | Normal Cognition |
| ast | inhib | VEGFA | ITGB1 | 1.18E-07 | 0.06268756 | Normal Cognition |
| ast | mic | VEGFA | ITGB1 | 1.18E-07 | 0.06268756 | Normal Cognition |
| ast | oligo | VEGFA | ITGB1 | 1.18E-07 | 0.06268756 | Normal Cognition |
| ast | opc | VEGFA | ITGB1 | 1.18E-07 | 0.06268756 | Normal Cognition |
| ast | ast | VEGFA | SIRPA | 4.72E-07 | 0.06865381 | Normal Cognition |
| ast | excit | VEGFA | SIRPA | 9.44E-07 | 0.07208409 | Normal Cognition |
| ast | inhib | VEGFA | SIRPA | 3.54E-07 | 0.06732413 | Normal Cognition |
| ast | mic | VEGFA | SIRPA | 1.18E-07 | 0.06268756 | Normal Cognition |
| ast | oligo | VEGFA | SIRPA | 1.18E-07 | 0.06268756 | Normal Cognition |
| ast | opc | VEGFA | SIRPA | 1.18E-07 | 0.06268756 | Normal Cognition |
| ast | excit | VEGFA | GRIN2B | 3.42E-06 | 0.07946072 | Normal Cognition |
| ast | inhib | VEGFA | GRIN2B | 3.90E-06 | 0.08028502 | Normal Cognition |
| ast | opc | VEGFA | GRIN2B | 4.72E-07 | 0.06865381 | Normal Cognition |
| ast | endo | VEGFA | FLT1 | 1.14E-06 | 0.07307029 | AD |
| ast | ast | VEGFA | ITGAV | 1.90E-07 | 0.06461113 | AD |
| ast | endo | VEGFA | ITGAV | 4.75E-08 | 0.05929965 | AD |
| ast | excit | VEGFA | ITGAV | 4.75E-08 | 0.05929965 | AD |
| ast | inhib | VEGFA | ITGAV | 4.75E-08 | 0.05929965 | AD |
| ast | mic | VEGFA | ITGAV | 4.75E-08 | 0.05929965 | AD |
| ast | opc | VEGFA | ITGAV | 1.90E-07 | 0.06461113 | AD |
| ast | excit | VEGFA | TYRO3 | 1.90E-07 | 0.06461113 | AD |
| ast | oligo | VEGFA | TYRO3 | 4.75E-08 | 0.05929965 | AD |
| ast | opc | VEGFA | TYRO3 | 4.75E-08 | 0.05929965 | AD |
| ast | ast | VEGFA | GPC1 | 4.75E-08 | 0.05929965 | AD |
| ast | excit | VEGFA | GPC1 | 1.42E-07 | 0.06343208 | AD |
| ast | inhib | VEGFA | GPC1 | 4.75E-08 | 0.05929965 | AD |
| ast | opc | VEGFA | GPC1 | 4.75E-08 | 0.05929965 | AD |
| ast | ast | VEGFA | ITGA9 | 4.75E-08 | 0.05929965 | AD |
| ast | excit | VEGFA | ITGA9 | 1.90E-07 | 0.06461113 | AD |
| ast | inhib | VEGFA | ITGA9 | 4.75E-08 | 0.05929965 | AD |
| ast | mic | VEGFA | ITGA9 | 4.75E-08 | 0.05929965 | AD |
| ast | opc | VEGFA | ITGA9 | 4.75E-08 | 0.05929965 | AD |

| | | | | | | |
|---|---|---|---|---|---|---|
| ast | ast | VEGFA | NRP1 | 4.75E-08 | 0.05929965 | AD |
| ast | endo | VEGFA | NRP1 | 4.75E-08 | 0.05929965 | AD |
| ast | excit | VEGFA | NRP1 | 4.75E-08 | 0.05929965 | AD |
| ast | mic | VEGFA | NRP1 | 4.75E-08 | 0.05929965 | AD |
| ast | ast | VEGFA | EGFR | 3.32E-07 | 0.06703493 | AD |
| ast | endo | VEGFA | EGFR | 1.90E-07 | 0.06461113 | AD |
| ast | inhib | VEGFA | EGFR | 1.42E-07 | 0.06343208 | AD |
| ast | opc | VEGFA | EGFR | 1.90E-07 | 0.06461113 | AD |
| ast | endo | VEGFA | FLT1 | 1.14E-06 | 0.07307029 | AD |
| ast | endo | VEGFA | NRP2 | 4.75E-08 | 0.05929965 | AD |
| ast | excit | VEGFA | EPHB2 | 4.75E-08 | 0.05929965 | AD |
| ast | inhib | VEGFA | EPHB2 | 4.75E-08 | 0.05929965 | AD |
| ast | ast | VEGFA | ITGB1 | 4.75E-08 | 0.05929965 | AD |
| ast | endo | VEGFA | ITGB1 | 1.90E-07 | 0.06461113 | AD |
| ast | excit | VEGFA | ITGB1 | 4.75E-08 | 0.05929965 | AD |
| ast | inhib | VEGFA | ITGB1 | 4.75E-08 | 0.05929965 | AD |
| ast | mic | VEGFA | ITGB1 | 4.75E-08 | 0.05929965 | AD |
| ast | oligo | VEGFA | ITGB1 | 4.75E-08 | 0.05929965 | AD |
| ast | opc | VEGFA | ITGB1 | 4.75E-08 | 0.05929965 | AD |
| ast | ast | VEGFA | SIRPA | 1.90E-07 | 0.06461113 | AD |
| ast | excit | VEGFA | SIRPA | 3.80E-07 | 0.0676404 | AD |
| ast | inhib | VEGFA | SIRPA | 1.42E-07 | 0.06343208 | AD |
| ast | mic | VEGFA | SIRPA | 4.75E-08 | 0.05929965 | AD |
| ast | oligo | VEGFA | SIRPA | 4.75E-08 | 0.05929965 | AD |
| ast | opc | VEGFA | SIRPA | 4.75E-08 | 0.05929965 | AD |
| ast | excit | VEGFA | GRIN2B | 1.38E-06 | 0.07409487 | AD |
| ast | inhib | VEGFA | GRIN2B | 1.57E-06 | 0.07481111 | AD |
| ast | opc | VEGFA | GRIN2B | 1.90E-07 | 0.06461113 | AD |

*Adjusted Probability = -1/log(prob.original)
